# Stem Cell Divisions, Driver Mutations, and Carcinogenesis in Purebred Dogs

**DOI:** 10.64898/2026.04.14.718346

**Authors:** Jack da Silva

## Abstract

Most cancers are initiated by mutations that inactivate tumour-suppressor genes or activate oncogenes. Fitting a multistage model of carcinogenesis to the increase in cancer mortality with breed-specific size and lifespan in dogs has predicted that four somatic driver mutations are typically required to initiate cancer. This result is reconsidered here because it depends on the relationship between the number of at-risk cells and breed weight. Using a power function for this relationship results in higher quality models that support single driver mutations activating oncogenes. In addition, parameter estimates suggest that somatic mutation rates increase with weight, likely because of reduced investment in somatic maintenance. Regression of cancer mortality on body weight and lifespan shows that 56% of cancers in dogs are the result of mutations arising from somatic cell division, compared to 66% in humans. A further 7% of cancers may be due to inherited recessive mutations deactivating tumour suppressor genes, as indicated by the relationship between cancer mortality and breed germline homozygosity. Some of the remaining unexplained variation in cancer mortality may be explained by germline mutations underlying breed predispositions to specific cancers. Contrasting results with humans provide novel insights into the dynamics of carcinogenesis.

## Introduction

Most cancers are initiated by driver mutations that inactivate tumour-suppressor genes or activate oncogenes [1-3]. Some of these mutations may be inherited, but germline mutations for cancer are overwhelming recessive [4], likely as a result of strong selection against rare dominant mutations of large effect. Therefore, driver mutations often arise in somatic cells [5]. In humans, it is estimated that 66% of cancers are due to mutations arising during stem-cell division, 29% are due to somatic mutations caused by environmental factors, and 5% are inherited [6, 7]. This effect of stem cell division was estimated from the correlation between the total number of stem cell divisions over an individual’s lifetime for a given tissue and the lifetime risk of cancer in that tissue across 31 tissues/cancers. The main conclusion was that the probability of driver mutations arising increases with the number stem cell divisions. Thus, larger tissues, larger bodies, or longer-lived individuals should be more susceptible to cancer. This is consistent with the observation that cancer mortality increases with height in humans [8-13] and with weight and lifespan in breeds of dog [14-18].

The increase in cancer mortality with size and lifespan across dog breeds may also be used to infer the role of stem-cell division in carcinogenesis. This approach has the advantage over comparison across tissues in that adaptive cancer suppression may vary among tissues with different susceptibilities to cancer, complicating the relationship between the number of stem-cell divisions and the risk of cancer [19-21], whereas selection for cancer suppression is expected to be weak within dog breeds [22]. Similarly, comparisons across human individuals avoids the confounding effects of among-tissue variation in cancer suppression [8]. Using this approach, Nunney [22] estimated that four somatic driver mutations are typically responsible for initiating carcinogenesis in dogs. This was accomplished by fitting a multistage (multiple driver mutation) model of carcinogenesis to data on cancer mortality, breed-specific weight, and breed-specific lifespan. A simplified version of the model assumes that a given cancer *i* is rare and estimates the prevalence of the cancer, *p*_*i*_, by age *T* as [23]:

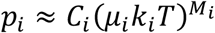

where *C* is the number of at-risk cells for the cancer, *k* is their rate of cell division, *μ* is the somatic mutation rate for driver mutations, and *M* is the number of driver mutations required to initiate the cancer. This equation gives the probability of *M* mutations accumulating in a single cell. Then, assuming *C* is proportional to body size, variation in the lifetime risk of cancer, body size, and lifespan across dog breeds may be used to estimate values for *M*_*i*_ from this equation. Nunney [22] did this using a modified version of the model that allows *C, μ*, and *k* to vary among cancer types with *M* driver mutations while fitting the model to the variation in *C* and *T* among breeds.

However, the results of Nunney [22] are shown here to depend on the assumption that *C* is a linear function of body weight, *W*, such that *C* = *fW*, where *f* is a constant of proportionality. Assuming an allometric (power) function, where *C* = *W*^*f*^, and using the highest quality data available on cancer mortality rates and lifespans for dog breeds, it is shown that models with improved fit to the data support single somatic driver mutations arising during stem-cell division, that is, a “one-hit” initiation via oncogenes [24]. Realistic interpretation of model parameters requires the assumption that the driver mutation rate increases with weight, consistent with the level of somatic maintenance declining with weight. This analysis indicates that 56% of cancer deaths in dogs may be explained by mutations arising during stem cell division, similar to the proportion in humans. The contribution of heritable mutations is also inferred from the relationship between cancer mortality and breed germline homozygosity. Some of the remaining unexplained variation in cancer mortality among breeds may be explained by germline mutations that underly breed predispositions to specific cancers. The analysis of cancer mortality in dog breeds may further our understanding of carcinogenesis in humans.

## Methods

### Data

Data on the breed-specific probability of dying from any cancer are from 81 breeds, each with 100 or more reported deaths, extracted from the Veterinarian Medical Database [16]. This dataset contains the most accurate data on cancer mortality available. Other large datasets rely on owner-reported causes of death, which likely underestimate cancer mortality [14, 15]. Adult body masses are the American Kennel Club (www.akc.org) breed standards for most breeds, with body masses for a few breeds from other sources [14]. Breed-specific individual lifespans are from Kraus, Snyder-Mackler [14], who estimated mean lifespans from owner-reported ages at death for breeds with at least 80 reported deaths. They used only 1988-2002 birth cohorts to avoid underestimating lifespans due to the inclusion of incomplete birth cohorts (right-censoring). Dogs that died due to extrinsic causes were excluded to provide an estimate of potential lifespan. Breed-specific median SNP heterozygosity is from a dataset for breeds with at least 30 genotyped individuals [25]. In the final dataset, there are 57 breeds with complete data on the probability of dying from cancer, body weight, lifespan, and heterozygosity.

### Analysis

Statistical analyses were conducted using the R language and environment for statistical computing [26]. Phylogenetic generalized least squares analyses [27], used to control for phylogenetic non-independence of breeds [28], showed no phylogenetic signal (*λ* = 0) in regressions of the probability of dying from cancer on any independent variable. Therefore, all regressions were conducted using either Levenberg-Marquardt nonlinear or ordinary linear least squares. The Akaike information criterion (AIC) was used in model selection, with values more than 2 units lower indicating significant improvement in model quality [29].

## Results

### Fitting the Multistage Model

Nunney [22] fit a modified multistage model of carcinogenesis to breed cancer mortality data. In this model, the probability of dying from cancer *i*, with *M*_*i*_ driver mutations, is a function of body weight, *W*, and lifespan, *T*:

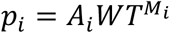

where *A*_*i*_ is an estimated parameter that includes the remaining model parameters: 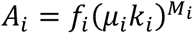, where *f*_*i*_ = *C*_*i*_⁄*W*. Note that this assumes *C*, the number of at-risk cells, is a linear function of *W*. Then, the overall probability of dying for breed *j* is

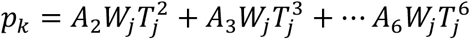

Each term in the equation (*e*.*g*., 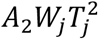) represents all cancers with *M* driver mutations (*e*.*g*., *M* = 2). The terms in the model span *M* = 2-6, reflecting plausible numbers of driver mutations. Forward stepwise nonlinear least squares regression was used to find the best combination of terms that describe the data. In the analysis, Nunney predicted *T* from a linear regression of lifespan on mean *W* for breed size classes using other data. This was done in an attempt to remove the potential dependence of lifespan on the risk of cancer. The model with the lowest AIC contains a single term, for *M* = 4 driver mutations: *p* = *A*_4_*WT*^4^ [22]. This result was confirmed using the data described in Methods.

However, a better fit to the data is obtained if *C* is assumed to be a power function of 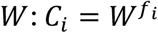. Now, the parameter *f*_*i*_ must be estimated separately and *A*_*i*_ is defined as 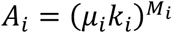:

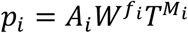

Nevertheless, forward stepwise regression shows that this model provides a significantly better fit to the data (lower AIC) if *M* = 1 or *M* = 2, indicating that one or two driver mutations explain the data most parsimoniously (see “*T* estimated from *W*” in Table 1; Fig. 1).

**Table 1.** Comparisons of modified evolutionary multistage models. The number of driver mutations, *M*, is specified for each model; *A* and *f* were estimated using Levenberg-Marquardt nonlinear least squares regression. Nunney’s model (*A*_4_*WT*^4^) is shown for reference; other models have lower or approximately equal AIC values for each method of estimating *T*. Models with significantly lowest AIC values for each method are in bold font.

| Model | $M$ | Estimate | | $df$ | AIC |
| --- | --- | --- | --- | --- | --- |
| | | $A$ | $f$ | | |
| $T$ estimated from $W$ | | | | | |
| $A_4WT^4$ | 4 | $7.99 \times 10^{-7}$ | — | 2 | −153.28 |
| $A_1W^{f_1}T^1$ | 1 | $7.05 \times 10^{-3}$ | 0.41 | 3 | −163.31 |
| $A_2W^{f_2}T^2$ | 2 | $3.94 \times 10^{-4}$ | 0.57 | 3 | −164.39 |
| $A_3W^{f_3}T^3$ | 3 | $2.22 \times 10^{-5}$ | 0.71 | 3 | −160.77 |
| $A_4W^{f_4}T^4$ | 4 | $1.25 \times 10^{-6}$ | 0.86 | 3 | −154.46 |
| $T$ is breed-specific lifespan | | | | | |
| $A_1WT^1$ | 1 | $1.01 \times 10^{-3}$ | — | 2 | −74.29 |
| $A_2WT^2$ | 2 | $1.02 \times 10^{-4}$ | — | 2 | −87.37 |
| $A_3WT^3$ | 3 | $9.68 \times 10^{-6}$ | — | 2 | −86.04 |
| $A_4WT^4$ | 4 | $8.74 \times 10^{-7}$ | — | 2 | −75.86 |
| $A_1W^{f_1}T^1$ | 1 | $7.24 \times 10^{-3}$ | 0.43 | 3 | −115.99 |
| $A_2W^{f_2}T^2$ | 2 | $4.11 \times 10^{-4}$ | 0.59 | 3 | −106.24 |
| $A_3W^{f_3}T^3$ | 3 | $2.36 \times 10^{-5}$ | 0.73 | 3 | −90.59 |
| $A_4W^{f_4}T^4$ | 4 | $1.40 \times 10^{-6}$ | 0.85 | 3 | −75.16 |

**Figure 1.**
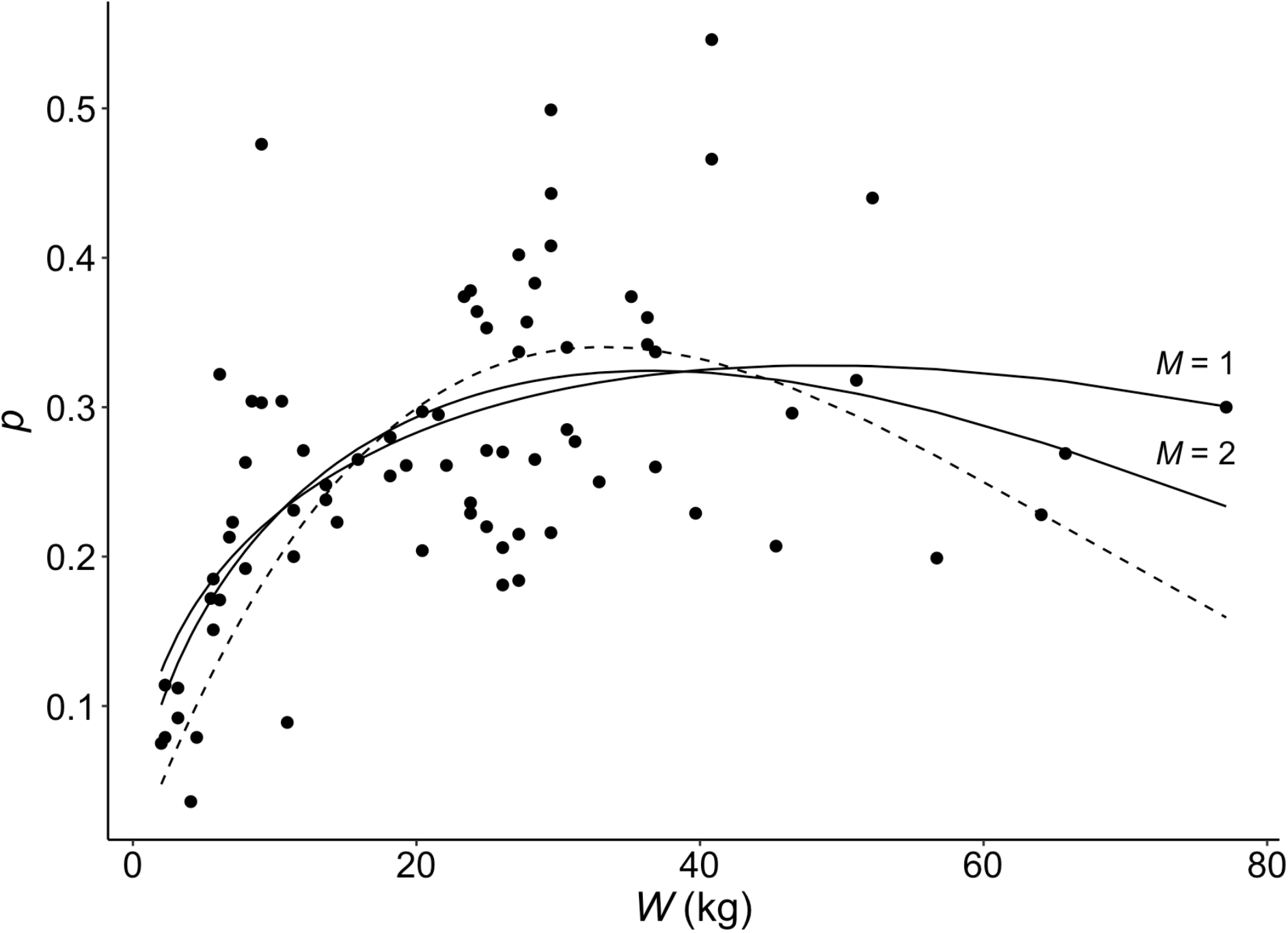
The probability of dying from cancer, *p*, as a function of weight, *W*, for Nunney’s model, *A*_4_*WT*^4^ (dashed line), and for models of the form *AW*^*f*^*T*^*M*^ that have significantly lower AIC values: *M* = 1 and *M* = 2 (solid lines). *T* is estimated from *W* for all models (see text).

The above analyses were conducted by first predicting *T* from a linear regression of lifespan on *W*, following Nunney [22], which makes *T* a function of *W* and thus not an independent variable in the analysis. Here, breed-specific lifespan is used for *T* instead. Now, Nunney’s model, with *M* = 4, no longer has the lowest AIC for models of that form (*C*_*i*_ = *f*_*i*_*W*); models of the same form but with *M* = 2 and *M* = 3 have significantly lower AIC values (see “*T* is breed-specific lifespan” in Table 1). However, a model with *C* as a power function of 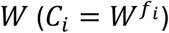 and *M* = 1 has the lowest AIC of all (Table 1). This suggests that cancers in purebred dogs are typically initiated by single driver mutations.

### Interpreting Model Parameters

The model with *C* as a power function of 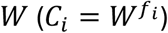 and *M* = 1 estimates *f* = 0.43 (see “*T* is breed-specific lifespan” in Table 1) and thus *C* = *W*^0.43^ at-risk cells. Therefore, 2.7, 3.6, and 5.8 at-risk cells are predicted for breeds weighing 10, 20, and 60 kg, respectively. These numbers of cells are much lower than the expected number of stem cells in most tissues. For example, in humans, which weigh approximately as much as a giant breed, there are on average ∼10^8^ stem cells per tissue, with a range of ∼10^5^ to ∼10^9^ stem cells across 31 tissues [6]. This large discrepancy between predicted and expected numbers of at-risk cells suggest that the parameters in the model are being interpreted incorrectly.

A solution to the above apparent misinterpretation of model parameters is to assume that as more resources are allocated to growth and reproduction with increasing breed weight, fewer resources are allocated to somatic maintenance, which may explain higher rates of senescence or higher base mortality rates in larger breeds [17, 30, 31]. Less investment in somatic maintenance may reduce the allocation of resources to the inhibition and repair of DNA damage, increasing mutation rates in larger breeds. Then, somatic mutation rates for driver mutations are expected to increase with weight. This hypothesis suggests that the model parameters should be interpreted as 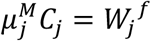 for breed *j* and *A* = *k*^*M*^. Now, with *M* = 1 and assuming *μ* = 10^−6^, a 60 kg breed is estimated to have *C* = 5.8 × 10^6^ at-risk cells, which is a more reasonable estimate. The relationship between the mutation rate and weight may explain why cancer mortality is a power function of weight rather than a linear function. However, now *k* = 7.24 × 10^−3^ cell divisions per year, which is much lower than a mean (range) of 13 (0-73) stem cell divisions per year across 31 tissues in humans [6]. This remaining problem with the interpretation of model parameters may be resolved by the use of a similar model that assumes single driver mutations.

### “One-Hit” Initiation via Oncogenes Model

The “one-hit” initiation via oncogenes model assumes a single (dominant) driver mutation activates an oncogene [24]:

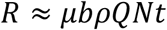

where *R* is the lifetime risk of cancer, *N* is the number of stem cells, *b* is their rate of division, *t* is organism age, *μ* is the probability of activating an oncogene multiplied by the number of oncogenes, *ρ* is the mean fixation probability of an oncogene mutation, and *Q* is the probability of cancer progression. Therefore, parameters are interpreted as *μN* = *W*^0.43^, *A* = *bρQ*, and thus *b* = 7.24 × 10^−3^⁄(*ρQ*) stem cell divisions per year. For small values of *ρ* and *Q*, say 0.02, then *b* = 18, which is reasonable.

### Sources of Mutations

#### Stem Cell Divisions

The one-hit model may be re-written in terms of weight, *W*, as for the multistage model:

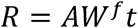

where again *A* = *bρQ* and *W*^*f*^ = *μN*. Then, taking the log of each side of the equation allows the use of linear ordinary least squares regression to estimate the parameters and to estimate the proportion of variance in log *R* explained by mutations arising from stem cell divisions:

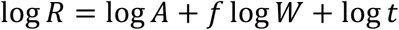

Regressing the log probability of dying from cancer on log breed weight and log lifespan gives the following equation (Table 2; Fig. 2; *R*^2^ = 0.56, *F*_2,54_ = 34.34; *P* = 2.384 × 10^−10^):

**Table 2.** Ordinary least squares regression of the log_10_ probability of dying from cancer, *R*, on log_10_ adult body mass, *W*, and log_10_ lifespan, *t*.

| Coefficient | Estimate | SE | $t$ value | $P$ |
| --- | --- | --- | --- | --- |
| Intercept | -2.6007 | 0.4688 | -5.547 | $8.97 \times 10^{-7}$ |
| $\log_{10} W$ | 0.5574 | 0.0712 | 7.835 | $1.83 \times 10^{-10}$ |
| $\log_{10} t$ | 1.2673 | 0.3915 | 3.237 | 0.0021 |

**Figure 2.**
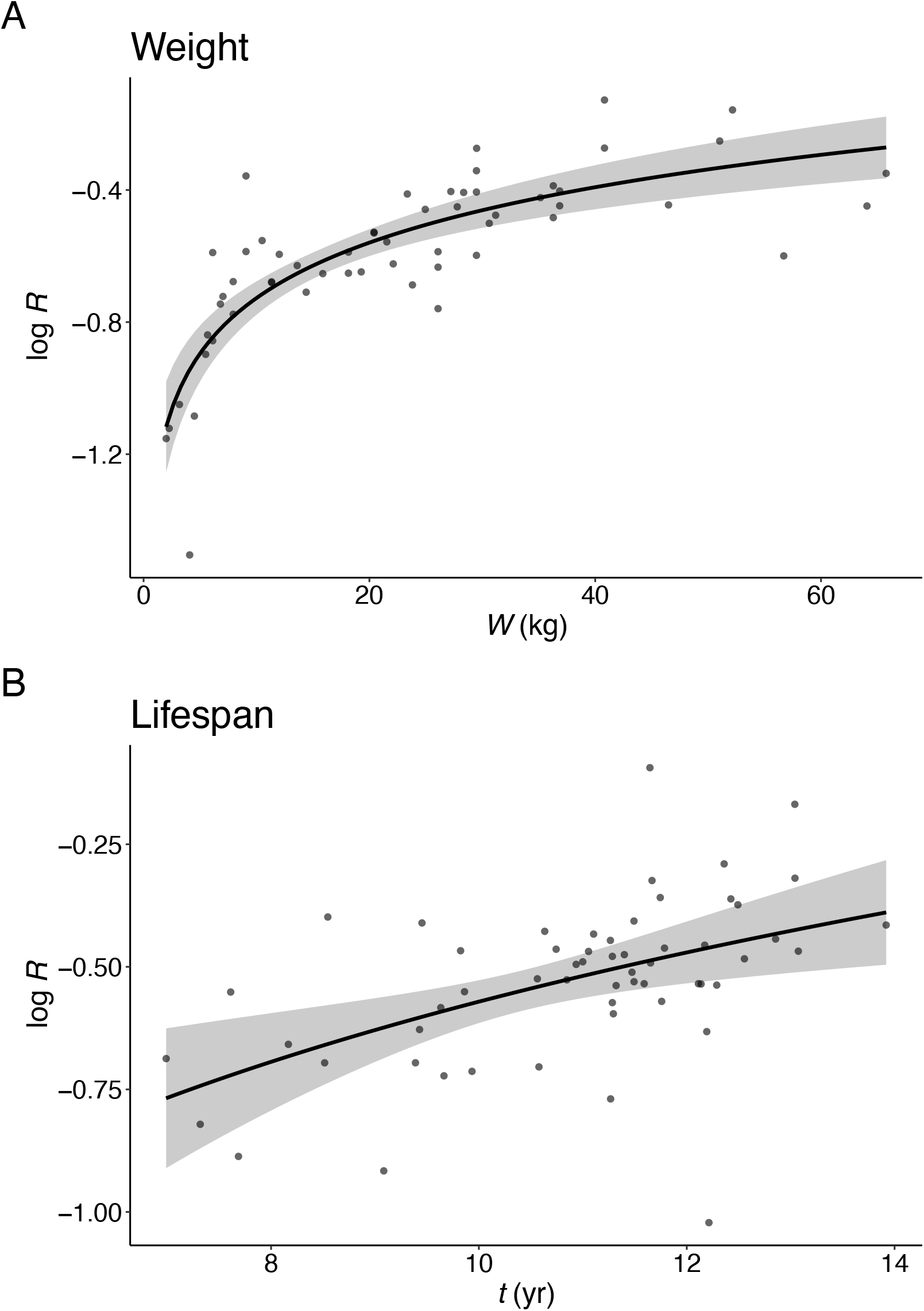
Partial residual plots from ordinary least squares multiple linear regression of the effects of (A) log weight, *W*, and (B) log lifespan, *t*, on the log probability of dying from cancer, *R*. 95% confidence bans of the regression lines are shown. Plots are shown with the *x*-axis on linear scale.

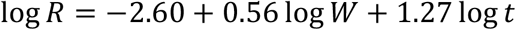

The slope of log *R* on log *t* is 1.27, which is not significantly different from 1 (*F*_54,55_ = 0.466, *P* = 0.4977), consistent with the one-hit model. The remaining parameters are interpreted as *N* = *W*^0.56^⁄*μ* stem cells and *b* = 2.51 × 10^−3^⁄(*ρQ*) stem cell divisions per year. The regression model also shows that 56% of the variance in log *R* is explained by log *W* and log *t*, which account for total stem cell divisions.

#### Germline Homozygosity

Intensive selective breeding often involves inbreeding, which increases germline homozygosity (1 – heterozygosity) [25, 32]. This may increase the probability that recessive mutations are found in the homozygous state, thus inactivating tumour suppressor genes [33]. Therefore, the probability of dying from cancer is predicted to increase with breed germline homozygosity after controlling for weight and lifespan to account for stem cell divisions. Adding breed homozygosity to the regression equation shows that log *R* increases with homozygosity and that homozygosity explains an additional 7% of variance in log *R*, supporting the hypothesis (Table 3; Fig. 3; *R*^2^ = 0.63, *F*_3,53_ = 29.93; *P* = 1.85 × 10^−11^).

**Table 3.**
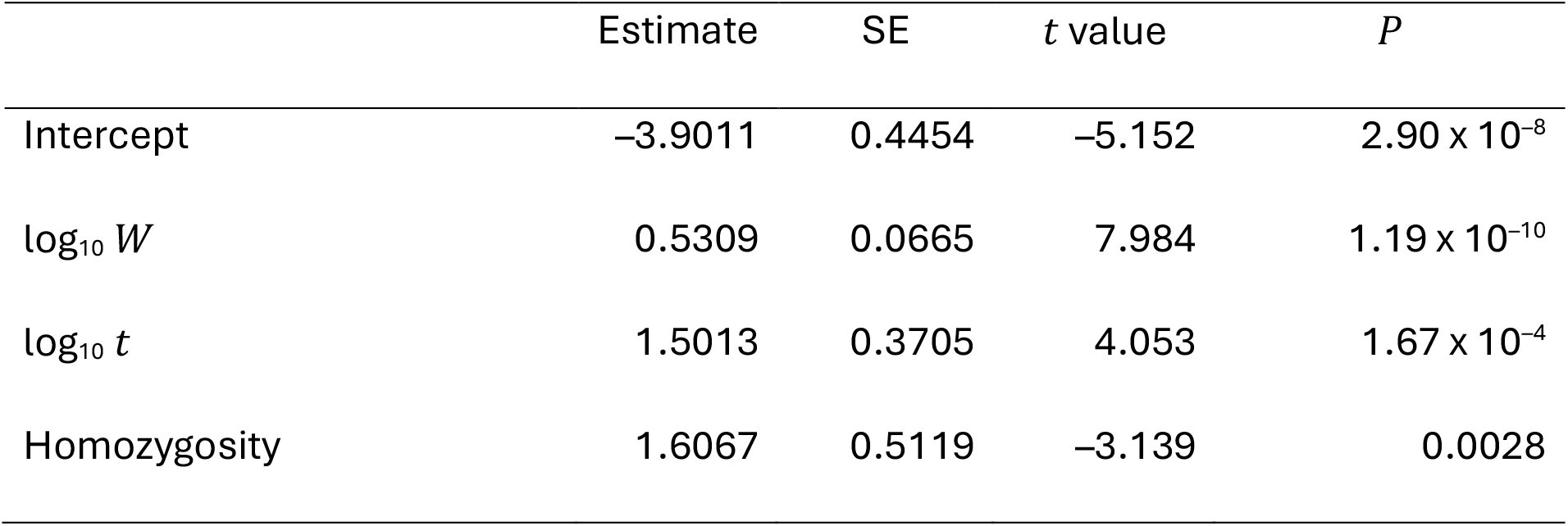
Ordinary least squares regression of the log_10_ probability of dying from cancer, *R*, on log_10_ adult body mass, *W*, log_10_ lifespan, *T*, and homozygosity.

**Figure 3.**
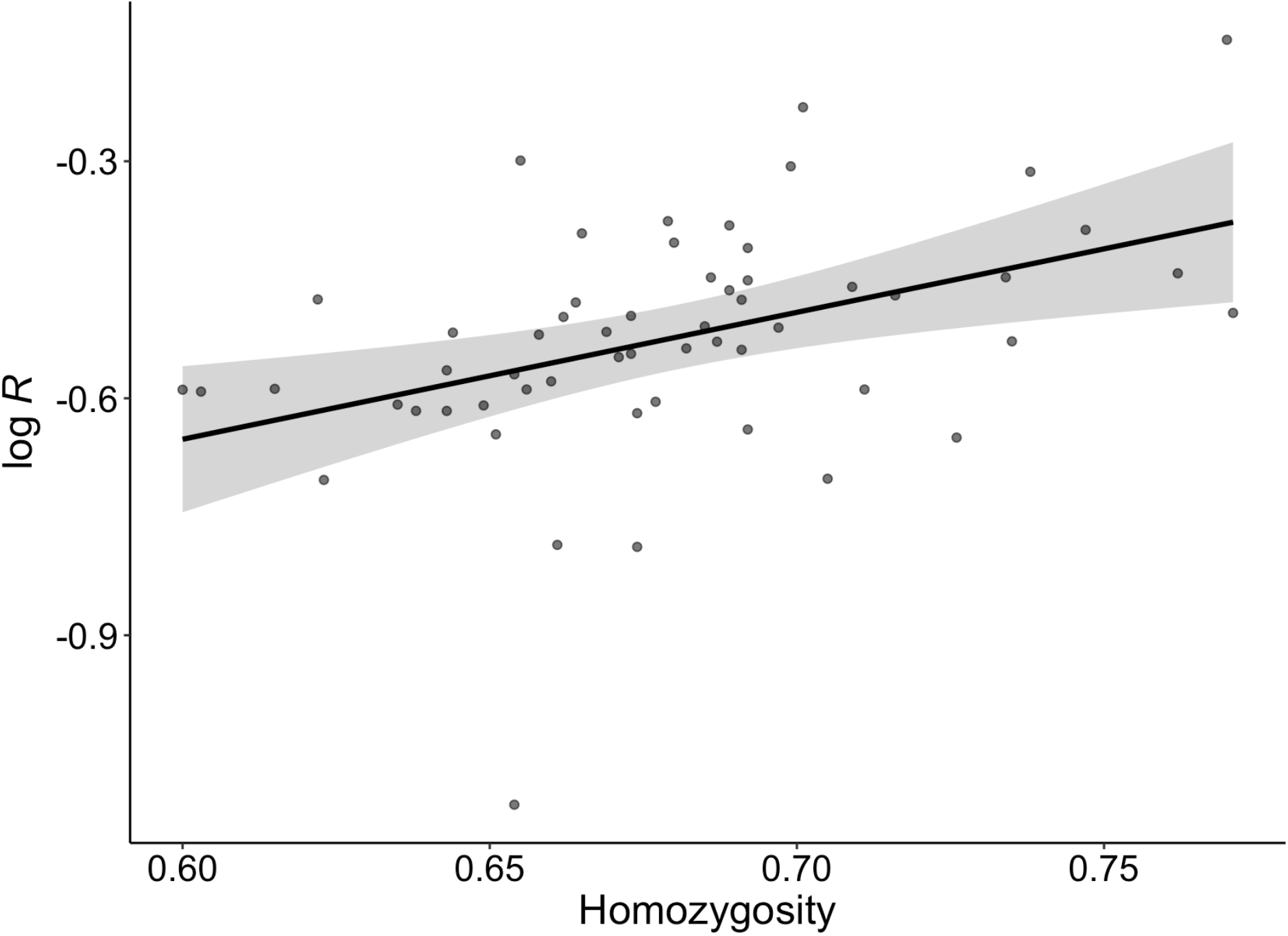
Partial residual plot of the effect of homozygosity on the log probability of dying from cancer, *R*, after accounting for weight and lifespan with ordinary least squares linear regression. The 95% confidence ban of the regression line is shown.

#### Breed Predispositions to Cancers

Strong selective breeding may result in breed predispositions to specific cancers because selected traits increase the risk of cancer or because of linkage between alleles at selected loci and cancer-causing mutations [34-36]. Therefore, some of the remaining unexplained cancer mortality in the regression model may be due to germline mutations that predispose some breeds to specific cancers. These may be dominant mutations of oncogenes or recessive mutations of tumour suppressor genes. This hypothesis predicts that breeds with a high residual (positive) probability of dying from cancer after controlling for weight, lifespan, and homozygosity are more likely to be predisposed to specific cancers than breeds with low residual (negative) cancer mortality. However, this prediction is difficult to test because the identification of breed predispositions may depend on the breed’s popularity or inclusion in a survey [35] and because there has been no large-scale, global quantitative survey of breed predispositions. Nevertheless, comparing breeds with residuals at and above the 90^th^ percentile to those with residuals below the 10^th^ percentile is suggestive. The types of tumours to which breeds are predisposed are from the broad survey of Dobson [35]. Surveys that are country specific were not used as these tend to have highly biased selections of breeds.

Six breeds have residuals at or above the 90^th^ percentile (0.15) (Table 4). Of these, the Scottish terrier and the Bernese mountain dog have highly breed-specific cancers. Scottish terriers have a 21-fold increased risk (relative to mixed-breed dogs) of invasive transitional cell carcinoma (invasive urothelial carcinoma), compared with a 2 to 7-fold increased risk for several other breeds [37]. No associated germline mutation has been identified in Scottish terriers, likely because the cancer is polygenic or because its causative alleles are approaching fixation, because they are in linkage with loci under selection, and thus cannot be detected by genome wide association studies [36]. In Shetland sheepdogs, this cancer is associated with a mutation in *NIPAL1*, which appears to affect the MAPK signalling pathway, known to be involved in tumour development and frequently dysregulated in both human and dog bladder cancers [38]. Bernese mountain dogs are predisposed to histiocytic sarcoma, with ∼15-25% of individuals developing the cancer [39], but this cancer is rare in most other breeds, with the notable exception of the flat-coated retriever, which was not in this dataset. An associated mutation in Bernese mountain dogs has been mapped to a haplotype spanning *MTAP* and part of *CDKN2A*, a tumour suppressor.

**Table 4.** Breeds with residual log_10_ *R* ≥ 90^th^ percentile and < 10^th^ percentile after controlling for weight, lifespan, and homozygosity. Tumour types are from Dobson [35].

| Residual | Breed | High Incidence Tumours |
| --- | --- | --- |
| $\geq 90^{\text{th}}$ Percentile | | |
| 0.2644 | Cairn Terrier |  |
| 0.2580 | Bernese Mountain Dog | histiocytic sarcoma |
| 0.2325 | Scottish Terrier | invasive transitional cell carcinoma; lymphoma; prostatic carcinoma; cutaneous melanoma |
| 0.1864 | Bullmastiff | mast cell tumours; lymphoma |
| 0.1562 | Bouvier des Flandres | lymphoma |
| 0.1492 | Pembroke Welsh Corgi |  |
| $< 10^{\text{th}}$ Percentile | | |
| -0.1354 | Greyhound | osteosarcoma |
| -0.2003 | Chow Chow | oral melanoma |
| -0.2187 | Pekingese |  |
| -0.2317 | Newfoundland |  |
| -0.2551 | Dalmatian |  |
| -0.5493 | Miniature Pinscher |  |

Six breeds have residuals below the 10^th^ percentile (–0.12) (Table 4). Of these, the greyhound is predisposed to osteosarcoma, which is common among large and giant breeds [35]. This cancer is likely a direct result of rapid bone growth due to selection for large size, which explains why the greyhound does not have high residual cancer mortality after controlling for weight. This explanation is supported by the fact that the closely related but much smaller whippet is not predisposed to osteosarcoma [35]. The development of osteosarcoma may also have an environmental component, in the form of stress imposed on bones, since racing dogs have a higher incidence of the cancer [35]. Osteosarcoma in greyhounds is associated with a 150 kb segment upstream from *CDKN2A/B* [40]. It is not clear whether oral melanoma in the chow chow is entirely explained by genetic risk or whether it is related to heavy pigmentation [35]. The risk of melanoma shows a higher increase with height in humans than other cancers, possibly due to a higher rate of cell division in skin epithelium [8]. The chow chow is a large ancient Asian breed, with only ∼13% modern European genetic admixture [41], and thus may also have had more time at its present large size to evolve increased cancer suppression in response to a greater risk of cancer.

## Discussion

The increase in cancer mortality with body weight and lifespan across dog breeds supports single driver mutations arising via stem cell division, a “one-hit” initiation via oncogenes model [24]. In humans, recent large-scale genome sequencing of tumours from 35 cancer types shows that the median number of driver mutations per tumour ranges from one to six, with an overall median of two, and with ∼50% of tumours containing a median of one driver mutation [Extended Data Fig. 6 in 1]. Therefore, the best available data for humans is in rough accord with single driver mutations in dogs, which have a 10-fold higher rate of cancer than humans [34].

In contrast, Nunney [22] estimates four somatic driver mutations in dogs. This difference is based on the assumed relationship between the number of at-risk cells and body weight. Nunney assumed a linear relationship, which is reasonable if each cell weighs the same. However, an allometric relationship results in significantly better model quality (significantly lower AIC). Furthermore, plausible estimates of numbers of at-risk cells require the assumption that the driver mutation rate increases with weight. That is, that larger breeds invest less in somatic maintenance and therefore experience higher somatic mutation rates. This assumption is supported by higher rates of senescence or base mortality in larger breeds, which is consistent with more resources being diverted to growth and reproduction from somatic maintenance in larger breeds [17, 30, 31].

In humans, it has been estimated that 66% of cancers are caused by mutations arising during stem cell division, 29% by somatic mutations due to environmental factors, and 5% by inherited mutations [6, 7]. In the present study, 56% of cancers are estimated to be caused by mutations arising from stem cell division in dogs. Recent strong selective breeding may have increased the proportion of inherited cancer-causing mutations in dogs [18, 34, 36]. A high proportion of inherited mutations is supported by a 10-fold higher rate of cancer in dogs compared to humans [34], even though dogs are smaller and shorter-lived, and by breed-specific dispositions to specific cancers [34-36].

Intensive selective breeding, which often involves inbreeding, is expected to increase germline homozygosity [25, 32]. This may result in increased homozygosity of recessive mutations that inactivate tumour suppressor genes [33]. Thus, 7% of cancers are inferred to be caused by recessive germline mutations from the relationship between cancer mortality and homozygosity. In humans, the vast majority of germline mutations that cause cancer are recessive, whereas the opposite is true for somatic mutations [4]. This difference between germline and somatic mutations is easily explained by the fact that dominant mutations are under strong selection when rare and thus efficiently removed from the germline. Neither Kraus, Snyder-Mackler [14] nor Nunney [22] found a significant effect of inbreeding or heterozygosity on cancer mortality in dogs. Kraus, Snyder-Mackler [14], using the same heterozygosity data analysed here, attribute the lack of an effect partly to the low quality of their data on causes of death, which were owner-reported. In the present study, the probability of dying from cancer was calculated from autopsy reports produced by qualified veterinarians [16].

Germline mutations underlying breed predispositions to specific cancers may explain some of the remaining unexplained variance in cancer mortality. These mutations may be breeder-selected because they also affect desirable traits or are inadvertently selected because they happen to be linked to selected alleles [35, 36]. Such mutations are expected to explain some of the excess cancer mortality after controlling for the effects of stem cell division (weight and lifespan) and inbreeding (germline homozygosity). Of the six breeds analysed here with residual cancer mortality ≥ the 90^th^ percentile, two have highly breed-specific predispositions to specific cancers. In one of these, the Bernese mountain dog, a mutation associated with histiocytic sarcoma has been mapped to a haplotype spanning part of the *CDKN2A* tumour suppressor gene [39].

A one-hit model explains carcinogenesis in dogs. Thus, fewer driver mutations are required to initiate cancer in dogs than in humans, for which a median of two driver mutations are required. This is consistent with the higher rate of cancer in dogs even though they are smaller and shorter lived than humans. Realistic interpretation of model parameters requires the assumption that the driver mutation rate increases with weight, consistent with the level of somatic maintenance declining with weight. Dogs have a similar but lower proportion of cancers initiated by mutations arising during stem cell division compared to humans. The proportion of cancers due to inherited mutations, however, may be higher in dogs, as expected from intense selective breeding. This is reflected in increased cancer mortality with germline homozygosity and in breed predispositions to specific cancers. These contrasts with humans provide novel insights into the dynamics of carcinogenesis.

## Acknowledgements

This work was supported by the School of Biological Sciences, University of Adelaide.

## References

1. Kinnersley B., Sud A., Everall A., Cornish A.J., Chubb D., Culliford R., Gruber A.J., Lärkeryd A., Mitsopoulos C., Wedge D., et al. 2024 Analysis of 10,478 cancer genomes identifies candidate driver genes and opportunities for precision oncology. Nature Genetics 56(9), 1868–1877. (doi:10.1038/s41588-024-01785-9).

2. Jassim A., Rahrmann E.P., Simons B.D., Gilbertson R.J. 2023 Cancers make their own luck: theories of cancer origins. Nature Reviews Cancer 23(10), 710–724. (doi:10.1038/s41568-023-00602-5).

3. Vogelstein B., Papadopoulos N., Velculescu V.E., Zhou S.B., Diaz L., Kinzler K.W. 2013 Cancer Genome Landscapes. Science 339(6127), 1546–1558. (doi:10.1126/science.1235122).

4. Futreal P.A., Coin L., Marshall M., Down T., Hubbard T., Wooster R., Rahman N., Stratton M.R. 2004 A census of human cancer genes. Nature Reviews Cancer 4(3), 177–183. (doi:10.1038/nrc1299).

5. Frank S.A., Nowak M.A. 2004 Problems of somatic mutation and cancer. Bioessays 26(3), 291–299. (doi:10.1002/bies.20000).

6. Tomasetti C., Vogelstein B. 2015 Variation in cancer risk among tissues can be explained by the number of stem cell divisions. Science 347(6217), 78–81. (doi:10.1126/science.1260825).

7. Tomasetti C., Li L., Vogelstein B. 2017 Stem cell divisions, somatic mutations, cancer etiology, and cancer prevention. Science 355(6331), 1330–1334. (doi:doi:10.1126/science.aaf9011).

8. Nunney L. 2018 Size matters: height, cell number and a person’s risk of cancer. Proceedings of the Royal Society B: Biological Sciences 285(1889).

9. Choi Y.J., Lee D.H., Han K.-D., Yoon H., Shin C.M., Park Y.S., Kim N. 2019 Adult height in relation to risk of cancer in a cohort of 22,809,722 Korean adults. British Journal of Cancer 120(6), 668–674. (doi:10.1038/s41416-018-0371-8).

10. Green J., Cairns B.J., Casabonne D., Wright F.L., Reeves G., Beral V. 2011 Height and cancer incidence in the Million Women Study: prospective cohort, and meta-analysis of prospective studies of height and total cancer risk. Lancet Oncology 12(8), 785–794.

11. Kabat G.C., Kim M.Y., Hollenbeck A.R., Rohan T.E. 2014 Attained height, sex, and risk of cancer at different anatomic sites in the NIH-AARP Diet and Health Study. Cancer Causes & Control 25(12), 1697–1706. (doi:10.1007/s10552-014-0476-1).

12. Sung J., Song Y.M., Lawlor D.A., Smith G.D., Ebrahim S. 2009 Height and Site-specific Cancer Risk: A Cohort Study of a Korean Adult Population. American Journal of Epidemiology 170(1), 53–64. (doi:10.1093/aje/kwp088).

13. Wirén S., Häggström C., Ulmer H., Manjer J., Bjorge T., Nagel G., Johansen D., Hallmans G., Engeland A., Concin H., et al. 2014 Pooled cohort study on height and risk of cancer and cancer death. Cancer Causes & Control 25(2), 151–159. (doi:10.1007/s10552-013-0317-7).

14. Kraus C., Snyder-Mackler N., Promislow D.E.L. 2023 How size and genetic diversity shape lifespan across breeds of purebred dogs. GeroScience 45(2), 627–643. (doi:10.1007/s11357-022-00653-w).

15. Adams V.J., Evans K.M., Sampson J., Wood J.L.N. 2010 Methods and mortality results of a health survey of purebred dogs in the UK. Journal of Small Animal Practice 51(10), 512–524. (doi:10.1111/j.1748-5827.2010.00974.x).

16. Fleming J.M., Creevy K.E., Promislow D.E.L. 2011 Mortality in North American dogs from 1984 to 2004: An investigation into age-, size-, and breed-related causes of death. Journal of Veterinary Internal Medicine 25(2), 187–198. (doi:10.1111/j.1939-1676.2011.0695.x).

17. da Silva J., Cross B.J. 2023 Dog life spans and the evolution of aging. The American Naturalist 201(6), E000–E000. (doi:10.1086/724384).

18. Doherty A., Lopes I., Ford C.T., Monaco G., Guest P., de Magalhães J.P. 2020 A scan for genes associated with cancer mortality and longevity in pedigree dog breeds. Mammalian Genome 31(7), 215–227.

19. Nunney L., Muir B. 2015 Peto’s paradox and the hallmarks of cancer: constructing an evolutionary framework for understanding the incidence of cancer. Philosophical Transactions of the Royal Society B: Biological Sciences 370(1673), 20150161. (doi:doi:10.1098/rstb.2015.0161).

20. Noble R., Kaltz O., Nunney L., Hochberg M.E. 2016 Overestimating the Role of Environment in Cancers. Cancer Prevention Research 9(10), 773–776. (doi:10.1158/1940-6207.Capr-16-0126).

21. Noble R., Kaltz O., Hochberg M.E. 2015 Peto’s paradox and human cancers. Philosophical Transactions of the Royal Society B: Biological Sciences 370(1673), 20150104. (doi:doi:10.1098/rstb.2015.0104).

22. Nunney L. 2024 The effect of body size and inbreeding on cancer mortality in breeds of the domestic dog: a test of the multi-stage model of carcinogenesis. R Soc Open Sci 11(1), 231356. (doi:10.1098/rsos.231356).

23. Nunney L. 1999 Lineage selection and the evolution of multistage carcinogenesis. Proceedings of the Royal Society of London Series B: Biological Sciences 266(1418), 493–498. (doi:10.1098/rspb.1999.0664).

24. Nowak M.A., Waclaw B. 2017 Cancer genes, environment, and “bad luck”. Science 355(6331), 1266–1267. (doi:10.1126/science.aam9746).

25. Bannasch D., Famula T., Donner J., Anderson H., Honkanen L., Batcher K., Safra N., Thomasy S., Rebhun R. 2021 The effect of inbreeding, body size and morphology on health in dog breeds. Canine Medicine and Genetics 8(1), 1–9.

26. R Core Team. 2025 R: A Language and Environment for Statistical Computing. (Vienna, Austria, R Foundation for Statistical Computing.

27. Pagel M. 1999 Inferring the historical patterns of biological evolution. Nature 401(6756), 877–884.

28. Parker H.G., Dreger D.L., Rimbault M., Davis B.W., Mullen A.B., Carpintero-Ramirez G., Ostrander E.A. 2017 Genomic analyses reveal the influence of geographic origin, migration, and hybridization on modern dog breed development. Cell Reports 19(4), 697–708.

29. Akaike H. 1974 A new look at the statistical model identification. IEEE Transactions on Automatic Control 19(6), 716–723. (doi:10.1109/TAC.1974.1100705).

30. Kraus C., Pavard S., Promislow D.E.L. 2013 The size–life span trade-off decomposed: Why large dogs die young. The American Naturalist 181(4), 492–505. (doi:10.1086/669665).

31. Bargas-Galarraga I., Vila C., Gonzalez-Voyer A. 2023 High investment in reproduction is associated with reduced life span in dogs. American Naturalist 201(2), 163–174. (doi:10.1086/722531).

32. Lewis T.W., Abhayaratne B.M., Blott S.C. 2015 Trends in genetic diversity for all Kennel Club registered pedigree dog breeds. Canine Genetics and Epidemiology 2(1), 13. (doi:10.1186/s40575-015-0027-4).

33. Ujvari B., Klaassen M., Raven N., Russell T., Vittecoq M., Hamede R., Thomas F., Madsen T. 2018 Genetic diversity, inbreeding and cancer. Proceedings of the Royal Society B: Biological Sciences 285(1875), 20172589. (doi:doi:10.1098/rspb.2017.2589).

34. Schiffman J.D., Breen M. 2015 Comparative oncology: what dogs and other species can teach us about humans with cancer. Philosophical Transactions of the Royal Society B: Biological Sciences 370(1673), 20140231.

35. Dobson J.M. 2013 Breed-predispositions to cancer in pedigree dogs. ISRN Veterinary Science 2013.

36. Ostrander E.A., Dreger D.L., Evans J.M. 2019 Canine cancer genomics: lessons for canine and human health. Annual Review of Animal Biosciences 7(1), 449–472.

37. Knapp D.W., Ramos-Vara J.A., Moore G.E., Dhawan D., Bonney P.L., Young K.E. 2014 Urinary bladder cancer in dogs, a naturally occurring model for cancer biology and drug development. ILAR journal 55(1), 100–118.

38. Parker H.G., Harris A.C., Plassais J., Dhawan D., Kim E.M., Knapp D.W., Ostrander E.A. 2024 Genome-wide analyses reveals an association between invasive urothelial carcinoma in the Shetland sheepdog and NIPAL1. npj Precision Oncology 8(1), 112. (doi:10.1038/s41698-024-00591-0).

39. Shearin A.L., Hedan B., Cadieu E., Erich S.A., Schmidt E.V., Faden D.L., Cullen J., Abadie J., Kwon E.M., Gröne A., et al. 2012 The MTAP-CDKN2A locus confers susceptibility to a naturally occurring canine cancer. Cancer epidemiology, biomarkers & prevention : a publication of the American Association for Cancer Research, cosponsored by the American Society of Preventive Oncology 21(7), 1019–1027. (doi:10.1158/1055-9965.Epi-12-0190-t).

40. Karlsson E.K., Sigurdsson S., Ivansson E., Thomas R., Elvers I., Wright J., Howald C., Tonomura N., Perloski M., Swofford R., et al. 2013 Genome-wide analyses implicate 33 loci in heritable dog osteosarcoma, including regulatory variants near CDKN2A/B. Genome Biology 14(12). (doi:ARTN R132 10.1186/gb-2013-14-12-r132).

41. Bergström A., Frantz L., Schmidt R., Ersmark E., Lebrasseur O., Girdland-Flink L., Lin A.T., Storå J., Sjögren K.-G., Anthony D., et al. 2020 Origins and genetic legacy of prehistoric dogs. Science 370(6516), 557. (doi:10.1126/science.aba9572).

